# miR-218 in adolescence predicts and mediates vulnerability to stress

**DOI:** 10.1101/2020.06.08.140038

**Authors:** Angélica Torres-Berrío, Alice Morgunova, Michel Giroux, Santiago Cuesta, Eric J. Nestler, Cecilia Flores

## Abstract

Adolescence is a period of increased vulnerability to psychiatric disorders including depression. Discovering novel biomarkers to identify individuals who are at high risk is very much needed. Our previous work shows that the microRNA miR-218 mediates susceptibility to stress and depression in adulthood, by targeting the Netrin-1 guidance cue receptor gene *Dcc* (*<u>D</u>eleted in <u>c</u>olorectal <u>c</u>ancer*) in the medial prefrontal cortex (mPFC). Here we investigated whether miR-218 regulates *Dcc* expression in adolescence and could serve as an early predictor of lifetime stress vulnerability. miR-218 expression in the mPFC increases from early adolescence to adulthood and correlates negatively with *Dcc* levels. In blood, postnatal miR-218 expression parallels changes occurring in the mPFC. Notably, circulating miR-218 levels in adolescence associate with vulnerability to social defeat stress in adulthood, with high levels associated with social avoidance severity. Indeed, downregulation of miR-218 in the mPFC in adolescence promotes resilience to stress in adulthood, indicating that adolescent miR-218 expression may serve both as a marker of risk and as a target for early interventions.

## INTRODUCTION

Adolescence is a developmental period of high-risk for the onset of stress-related disorders, including Major Depressive Disorder (MDD) and anxiety(1–6). Depression is a debilitating psychiatric condition and is the most prevalent psychopathology emerging in adolescence with a lifetime prevalence of approximately 25%(2, 7, 8). The current diagnosis of MDD is based on a self-report of symptoms, such as constant feelings of sadness, lack of interest or pleasure, hopelessness or suicidal ideation(9). Many of these symptoms can be observed during early adolescence(7, 10) and are exacerbated by environmental factors, including a lifetime history of stress(11). In high risk individuals, MDD symptomatology in adolescence can persist into adulthood and is a predictor of the severity of depression later in life(7, 10). The prevalence of MDD is alarming, but given its high heterogeneity, there is still a lack of biological, objective factors – biomarkers – to identify vulnerable youth or to predict responses to a pharmacological treatment. Current intervention strategies are therefore suboptimal for a large percentage of patients(12).

In search of objective biomarkers, several longitudinal studies have been conducted using peripheral fluids, including blood, saliva, or urine, to correlate the levels of stress-related hormones (e.g. cortisol) or inflammatory markers (e.g. Interleukin 6 and C-reactive protein) with depressive symptoms and mood variations during adolescence(13–21). One limitation of these studies is the fact that depressive symptoms can be experienced in the absence of alterations in stress systems, including the hypothalamic–pituitary–adrenal (HPA) axis(22). The use of transcriptional profiles of mRNAs and non-coding RNAs in blood has been proposed to serve as a proxy of transcriptional activity in the brain(23–25), and potential adolescent biomarkers of vulnerability to depression have begun to emerge(26, 27). Several studies report altered levels of specific microRNAs (miRNA), small RNAs that repress gene expression, in the adult brain that are linked to MDD (see reviews by(23–25)) or can predict antidepressant response to pharmacological and behavioral interventions(23, 28, 29).

Our work identified miR-218 as a molecular switch of susceptibility versus resilience to stress-related disorders by a mechanism that involves the Netrin-1 guidance cue receptor, DCC (<u>d</u>eleted in <u>c</u>olorectal <u>c</u>ancer)(30, 31). DCC participates in the organization of neuronal circuitry across the lifespan and play a critical role in the maturation of the prefrontal cortex (PFC) during adolescence(32–34). Numerous genome-wide association studies (GWAS) have identified specific single-nucleotide polymorphisms (SNPs) within the *DCC* gene to be linked to MDD(35–43) (for review see:(44, 45)). We recently showed that miR-218 and DCC protein are coexpressed by PFC neurons and that miR-218 binds directly to the 3’-untranslated region (3’UTR) of *DCC* mRNA to repress its expression(30, 46). Adult subjects with MDD who died by suicide display reduced expression of miR-218 and increased expression of *DCC* mRNA in the PFC(32), with the levels of both transcripts correlating in a negative manner(31). These human findings were recapitulated in studies with adult mice susceptible to chronic social defeat stress (CSDS)(30, 31), a validated mouse model of stress-induced depression-like behaviors. Specifically, reducing levels of miR-218 in the medial PFC (mPFC) in adulthood increases vulnerability to social defeat and other stresses(31). In contrast, selective overexpression of miR-218 or downregulation of DCC itself in pyramidal neurons of the mPFC not only prevents depression-like behaviors(30, 31), but also protects against CSDS-induced morphological alterations(31).

A key discovery of our studies is that adult susceptible mice to CSDS also display reduced levels of miR-218 in blood, and in fact circulating expression correlates positively with social avoidance(31). Remarkably, experimentally-induced miR-218 downregulation in the adult mPFC causes a parallel reduction in circulating miR-218 levels, whereas viral-mediated upregulation of miR-218 selectively in adult mPFC pyramidal neurons leads to increased miR-218 levels in blood. These findings suggest that circulating miR-218 might serve as a sensor of miR-218/DCC changes occurring in the mPFC following adult exposure to chronic stress(31), providing information about effects of environmental risk and protective factors. However, whether circulating levels of miR-218 could serve as an early predictor biomarker of vulnerability to stress needs to be determined.

Given the important role of DCC receptors in the adolescent maturation of the mPFC(34, 44), we hypothesize that miR-218 determines *Dcc* gene expression in the mPFC during this sensitive developmental window and may mediate vulnerability to stress in adulthood. We also propose that circulating levels of miR-218 might serve as a developmental biomarker of depression-like behaviors in adulthood. To address these questions, we assessed whether the levels of miR-218 and *Dcc* mRNA in the mPFC vary across postnatal development and whether these variations can be detected in blood. We determined whether circulating miR-218 in adolescence predicts individual differences in vulnerability to CSDS in adulthood.

## METHODS AND MATERIALS

### Animals

All experimental procedures were performed in accordance with the guidelines of the Canadian Council of Animal Care and approved by the McGill University and Douglas Hospital Animal Care Committee.

#### C57BL/6 mice

Adolescent (PND 21 and PND 35) and adult (PD 75±15) male wild-type mice were bred in the animal facilities of the Douglas Hospital. Male pups were weaned at PND 21, group-housed (4 per cage) with same-sex littermates and given access to food and water *ad libitum*. All mice were kept on a 12-hour light/dark cycle (lights on 08:00), with experimental procedures being performed during the light period.

#### CD-1 mice

Male CD-1 retired breeder mice (>3 months old) obtained from Charles River Canada were used as aggressors in the chronic social defeat stress paradigm. CD-1 mice were single-housed throughout the study.

### Chronic Social Defeat Stress Paradigm (CSDS)

The CSDS protocol was conducted as in(30, 31) and consisted of a daily session in which an adult C57BL/6 experimental mouse was exposed to 5 min of physical aggression by a CD-1 mouse followed by overnight housing on a two compartment rat cage separated by a transparent and perforated central divider that allowed sensory but not physical contact between mice. The procedure was repeated for 10 consecutive days, in which C57BL/6 experimental mice were exposed to a new aggressor every day. Control C57BL/6 mice were housed in similar 2-compartment rat cages with a different littermate every day.

#### Social interaction test

Twenty-four hours after the last session of CSDS, C57BL/6 experimental mice were assessed in the social interaction test to measure their approach and/or avoidance behavior towards a social target as before(30, 31). Briefly, the social interaction test consisted of 2 sessions in which defeated and control mice were allowed to explore a squared-arena (42 cm x 42 cm) for a period of 2.5 minutes each session in the absence (session 1) or presence (session 2) of a novel CD-1 mouse. The time spent (in seconds) in the interaction zone and the corners was recorded during both sessions of the test. The social interaction ratio was calculated as the time spent in the interaction zone with the CD-1 aggressor present divided by the time spent in the interaction zone with the CD-1 aggressor absent. Defeated mice with a ratio 1 were classified as susceptible and a ratio≥1 were classified as resilient. Animal behavior was recorded with an overhead video camera for offline analysis using TopScanTM 2.0 software (Clever Systems Inc., Reston, VA, USA).

### Tissue Dissection and RNA Extraction

Wildtype male C57 mice were euthanized by decapitation at PND 21, PND35 and adult ages. Brains were removed and flash frozen with 2-methylbutane chilled in dry ice. Bilateral punches of the pregenual medial PFC, including prelimbic (PrL), and infralimbic (IL) subregions, were taken from 1mm coronal sections corresponding to plates 15-18 of the Paxinos & Franklin mouse atlas(47) and processed for gene expression experiments as previously(30). Total RNA and miRNA fractions were isolated from the mouse frozen tissue with the miRNeasy Micro Kit protocol (Qiagen, Toronto, ON, Canada). All RNA samples were determined to have 260/280 and 260/230 values ≥1.8, using the Nanodrop 1000 system (Thermo Fisher Scientific, Toronto, ON, Canada).

### Blood Collection

Blood samples were obtained before (submandibular blood) and after (trunk blood) exposure to the chronic social defeat stress paradigm, as described below:

#### Submandibular blood collection

Adolescent wildtype mice (PND 35) were restrained using the one-handed technique. A small puncture at the back of the jaw, slightly behind the hinge of the jawbone and toward the ear, was done with a 22 gauge needle. A maximum of 7.5% of the body weight was obtained in BD vacutainer tubes containing EDTA as anticoagulant (VWR, Ville Mont-Royal, Quebec). Adolescent mice were closely monitored for bleeding, or signs of pain, or distress, for one week after the procedure.

#### Trunk blood collection

Adult mice were decapitated 24 hours after the social interaction test, as described above. Blood from trunk was rapidly collected in BD vacutainer tubes containing EDTA as anticoagulant (VWR, Ville Mont-Royal, Quebec).

#### RNA extraction

Total RNA from whole blood, including miRNA fraction, was isolated using the Total RNA Purification Kit protocol (Norgen Biotek Corp., Thorold, ON, Canada), according to the manufacturer’s instructions (with minor modifications). All RNA samples were determined to have 260/280 and 260/230 values ≥1.8, using the Nanodrop 1000 system (Thermo Fisher Scientific, Toronto, ON, Canada).

### Quantitative Real Time PCR for Mouse Brain Tissue and Whole Blood

Reverse transcription for *Dcc* and *Robo-1* mRNA was performed as previously(46). Reverse transcription for miR-218 was performed using the TaqMan MicroRNA Reverse Transcription Kit and TaqMan probes (Applied Biosystems, QC, Canada). Real-time PCR was run in technical triplicates with an Applied Biosystems QuantStudio RT PCR system (Applied Biosystems, QC, Canada). The small nucleolar RNA (snoRNA) RNU6B was used as an endogenous control of miRNA measures. Expression levels of miR-218 were calculated using the Absolute Quantitation (AQ) standard curve method and normalized by RNU6B. The expression of miR-218 was relativized to (i.e., normalized to) the control group and presented as a fold change.

### Stereotaxic Surgery

Intracranial surgeries were performed under aseptic conditions as described previously(30, 31). Adolescent male (PND ~40) C57BL/6 wild-type mice were deeply anesthetised with isoflurane (5% for induction and 2% for maintenance) and placed in a stereotaxic apparatus. Bilateral microinfusions were done using stainless-steel infusion cannulae (33 gauge) into the mouse mPFC, including PrL and IL subregions, at the following coordinates: +2 mm (A/P), ±0.5 mm (M/L), and −2.7 mm (D/V) relative to Bregma. A locked nucleic acid (LNA) oligonucleotide with a sequence targeting miR-218 (Ant-miR-218) was used to downregulate the expression of miR-218 in vivo (Exiqon, Vedbaek, Denmark). A scrambled LNA oligonucleotide sequence was used as a control (Ant-scrambled). Ant-miR-218 or Ant-scrambled was dissolved in sterile PBS (Sigma-Aldrich, Oakville, ON, Canada) at a final concentration of 0.3nM and infused with a total volume of 0.5μl of Ant-miR-218 or Ant-scrambled over a 7-min period as in(31, 46). Mice exposed to CSDS, were left undisturbed until they reached adulthood, whereas mice used for validation were allowed to recover for 7 days prior to the sacrifice.

### Statistical Analysis

All values were represented as scatterplot with the mean ± s.e.m. Statistical analysis was performed using Graphpad Prism 6.0. A significance threshold of α <0.05 was used in all experiments. Statistical differences between two groups were analyzed with Student’s t-tests. Correlations were calculated using the Pearson correlation coefficient with two-tailed analysis. Otherwise one-way or two-way ANOVAs were performed, followed by Tukey’s multiple comparison or paired-t tests. No statistical methods were used to determine the sample sizes, but the number of experimental subjects is similar to sample sizes routinely used in our laboratory and in the field for similar experiments. All data were normally distributed and variance was similar between groups, supporting the use of parametric statistics.

## RESULTS

### Dynamic and opposite postnatal expression of miR-218 and *Dcc* mRNA in the mPFC

DCC is conspicuously expressed by cortical neurons throughout life, but levels decrease from adolescence onwards(48). To examine whether miR-218 regulates *Dcc* mRNA expression in the mPFC across postnatal life, we collected bilateral punches of the pregenual PrL and IL sub-regions of the mPFC of wildtype mice at three developmental ages: early adolescence (postnatal day, PND21), mid-adolescence (PND35), and adulthood (PND75), defined according to our previous work(49) **(FIGURE 1A)**. Using quantitative-PCR, we find that levels of miR-218 in the mPFC increase from early adolescence to adulthood **(FIGURE 1B)**, whereas the expression of *Dcc* mRNA decreases following the exact opposite pattern **(FIGURE 1C)**. Consistent with our previous findings(30, 46), levels of miR-218 and *Dcc* mRNA in the mPFC are inversely correlated **(FIGURE 1D)**, strongly suggesting that miR-218 controls *Dcc* gene expression in the mPFC throughout life.

**FIGURE 1.**
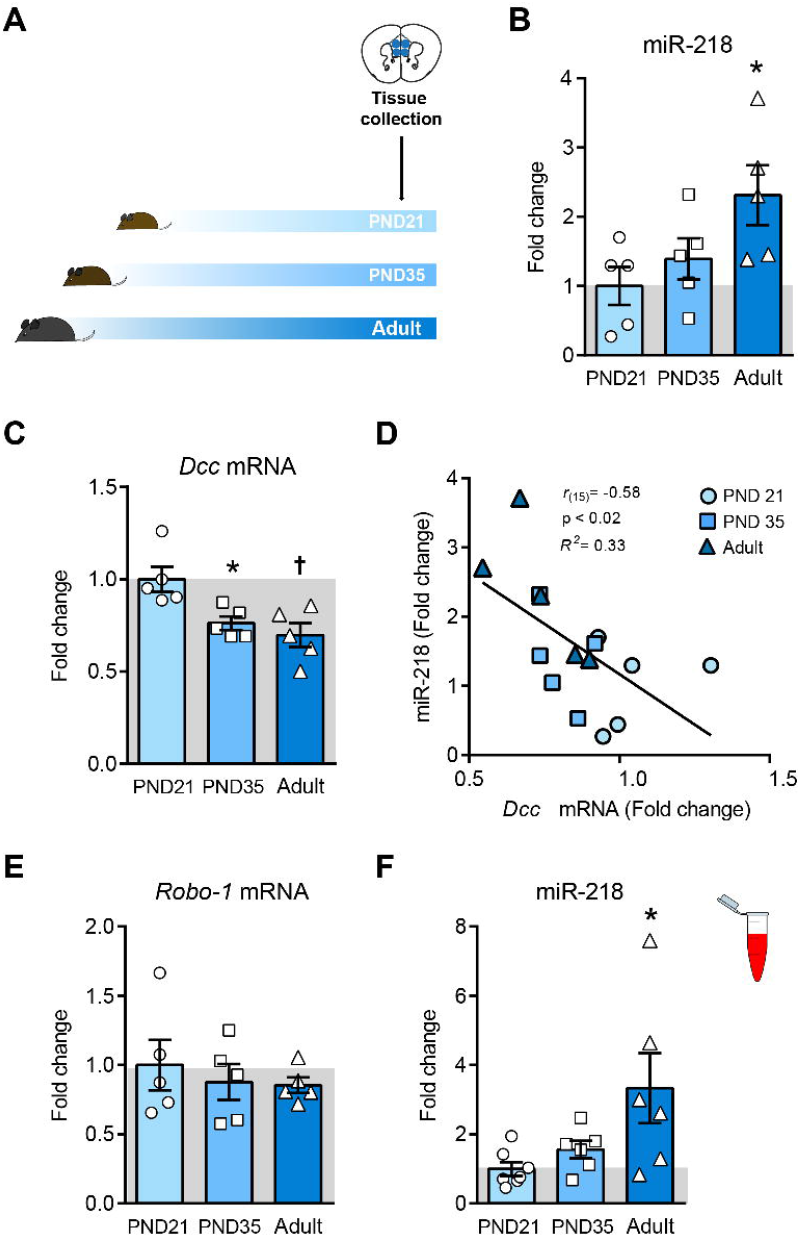
Developmental expression of miR-218 and *Dcc* and mRNA in the mPFC. **(a)** Timeline of experiment. **(b)** miR-218 increases from early adolescence (PND21) to adulthood (PND75±15). One-way ANOVA: F_(2,12)_ = 3.9; p<0.05; Tuckey test: *p□0.05; different from PND 21. **(c)** *Dcc* mRNA decreases in postnatal life. One-way ANOVA: F_(2,12)_ = 7.55; p<0.01; Tuckey test: *p□0.05; different from PND 21; †p□0.01; different from PND 35. **(d)** Negative correlation between PFC levels of *Dcc* mRNA and miR-218 across the lifespan. **(e)** The expression of *Robo-1* mRNA does not change from adolescence to adulthood. One-way ANOVA: F_(2,12)_ = 0.35; p=0.71. **(f)** miR-218 expression in blood increases across the lifespan. One-way ANOVA: F_(2,16)_= 4.341; p<0.05; Tuckey test: *p□0.05; different from PND 21.

In addition to *Dcc* mRNA, miR-218 also regulates the expression of *Robo-1* mRNA, which encodes for the receptor of the guidance cue, SLIT(50, 51). DCC and ROBO-1 form intracellular complexes that alter the attractant response to their ligand, Netrin-1(51). We measured *Robo-1* mRNA expression in the mPFC across the lifespan. Unlike *Dcc* mRNA levels, there are no significant changes in the expression of *Robo-1* mRNA from adolescence to adulthood **(FIGURE 1E**). Thus, while the developmental expression of *Dcc* in the mPFC appears to be controlled by opposite variations in miR-218 levels, *Robo1* expression is not sensitive to these changes. This finding suggests that miR-218 regulates certain guidance cue receptor genes to orchestrate the effects of environmental factors with high selectivity(45).

### Blood levels of miR-218 match its expression in the mPFC across postnatal development

Variations of miR-218 in the mPFC of adult mice are detected in peripheral blood, and circulating levels correlate with depression-like behaviors(31). To determine whether this is the case across postnatal life, we collected blood samples from a separate group of wildtype mice at PND21, PND35, and adulthood to measure circulating miR-218. Blood expression of miR-218 also increases from early adolescence to adulthood **(FIGURE 1F)**. This supports the notion that, across the lifespan, circulating levels of miR-218 mirror changes occurring in the mPFC.

### Circulating miR-218 in adolescence predicts susceptibility to CSDS in adulthood

miR-218 levels in blood of adult mice correlate with CSDS-induced social avoidance(31), suggesting a potential role for miR-218 as a peripheral biomarker of risk. Since adolescence is a developmental period of high risk for the onset of MDD(2), we measured blood levels of miR-218 in adolescence to determine whether they are associated with future susceptibility to depression-like behaviors induced by CSDS in adulthood. We collected blood samples of stress-naïve wildtype mice at PND35. We selected this developmental age because it represents a postnatal period of transition for miR-218 expression in both mPFC and blood **(FIGURE 1B,F)**. Adolescent mice were left undisturbed until they reached adulthood, when they underwent CSDS exposure **(FIGURE 2A)** or were assigned to the control group. Based on the time spent in the social interaction zone in the absence and presence of a novel social target (CD-1 mouse), we segregated socially-stressed mice into susceptible and resilient populations, based on the social interaction ratio as in(30, 31). By definition, CSDS significantly reduces social interaction in susceptible mice only **(FIGURE 2B)**.

**FIGURE 2.**
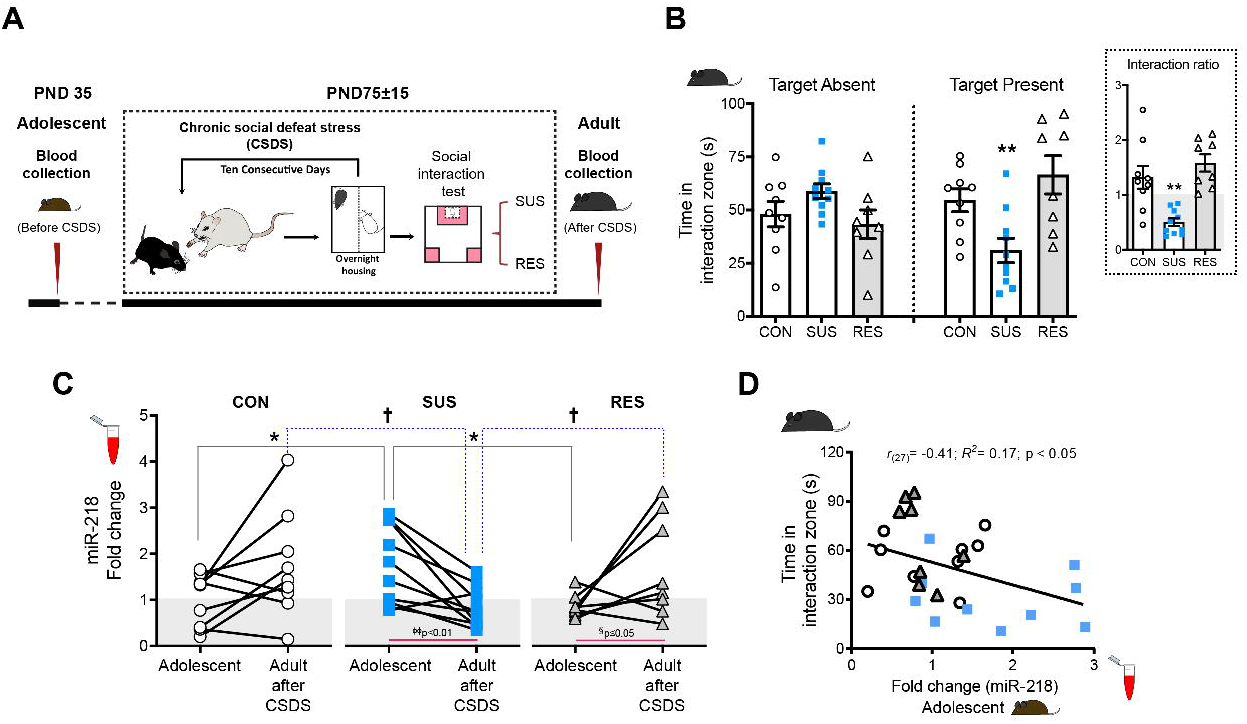
Levels of miR-218 in blood during adolescence correlate with susceptibility to CSDS in adulthood. **(a)** Timeline of blood collection and CSDS protocol. **(b)** Time in the interaction zone in control (CON=9), susceptible (SUS=10), resilient (RES=8) mice. Two-way ANOVA: significant stress by target interaction: F_(2,48)_= 9.44; p□0.001; stress: F(2,48)= 1.43; p= 0.2; session: F(1,48)= 0.01; p=0.9. Post hoc Tukey test: **p□0.01; different from CON and RES, in session with target present. Inset: Social interaction ratio: One-way ANOVA: F_(2,24)_= 14.02; p<0.0001; Tuckey test: **p□0.01; different from CON and RES. **(c)** Blood levels of miR-218 in adolescence and following CSDS in adulthood. All samples were segregated retrospectively into CON, SUS, and RES groups according to the social interaction ratio as in(30, 31). Expression was normalized to CON in Adolescent. Two-way ANOVA: Age by stress interaction: F_(2,48)_= 7.18; p<0.01; age: F_(1,48)_= 1.06; p= 0.31, stress: F_(2,48)_= 0.040; p= 0.96. Paired t-test: *p□0.05; SUS different from CON and RES in Adolescent. †p□0.05; SUS different from CON and RES in Adult after CSDS. **(d)** Negative correlation between adolescent levels of miR-218 in blood and the time in the interaction zone with target present.

We next used this adult phenotypic classification (control, susceptible and resilient) to assess group differences in circulating levels of miR-218 in blood samples obtained (*a*) in adolescence and (*b*) twenty-four hours after the social interaction test in adulthood **(FIGURE 2A)**. Analysis of blood samples in adolescence revealed that mice that become susceptible to CSDS display higher circulating miR-218 levels during this period, compared to levels observed in control and resilient groups **(FIGURE 2C)**. Consistent with the increase of miR-218 in circulating levels across development, analysis of blood samples after CSDS shows that adult control mice exhibit slightly higher levels of miR-218 in blood in comparison to adolescence **(FIGURE 1F)**. While resilient mice exhibit this same pattern of adolescence-to-adult increase, susceptible mice have *lower* levels of circulating miR-218 after CSDS **(FIGURE 2C)**. These results replicate our previous finding showing low levels of circulating miR-218 in adult mice susceptible to CSDS(31). These data also suggest that mice that exhibit “adult-like levels” of miR-218 in blood during adolescence are more vulnerable to the deleterious effects of CSDS in adulthood. Indeed, there is a significant *negative* correlation between circulating levels of miR-218 in adolescence and the time adult mice spend in the interaction zone with the social target (CD-1) present **(FIGURE 2D)**. To our knowledge, this is the first report showing that adolescent levels of a specific miRNA can provide information about prospective susceptibility versus resilience to CSDS in adulthood.

### Suppressing miR-218 in the mPFC in adolescence confers resilience to stress in adulthood

Because high levels of circulating miR-218 in adolescence correlates with CSDS-induced social avoidance in adulthood **(FIGURE 2D)**, we next examined whether reducing miR-218 expression in the mPFC of adolescent mice would influence susceptibility to adult exposure to CSDS. We used a nucleic acid (LNA)-modified antisense oligonucleotides (antagomir) that hybridize miR-218 (Ant-miR-218) to induce its degradation (**FIGURE 3A**). We used a scrambled sequence (Ant-Scrambled) as control. Adolescent mice were microinfused with either Ant-miR-218 or Ant-Scrambled into the mPFC and allowed to reach adulthood for CSDS exposure (**FIGURE 3B**).

**FIGURE 3.**
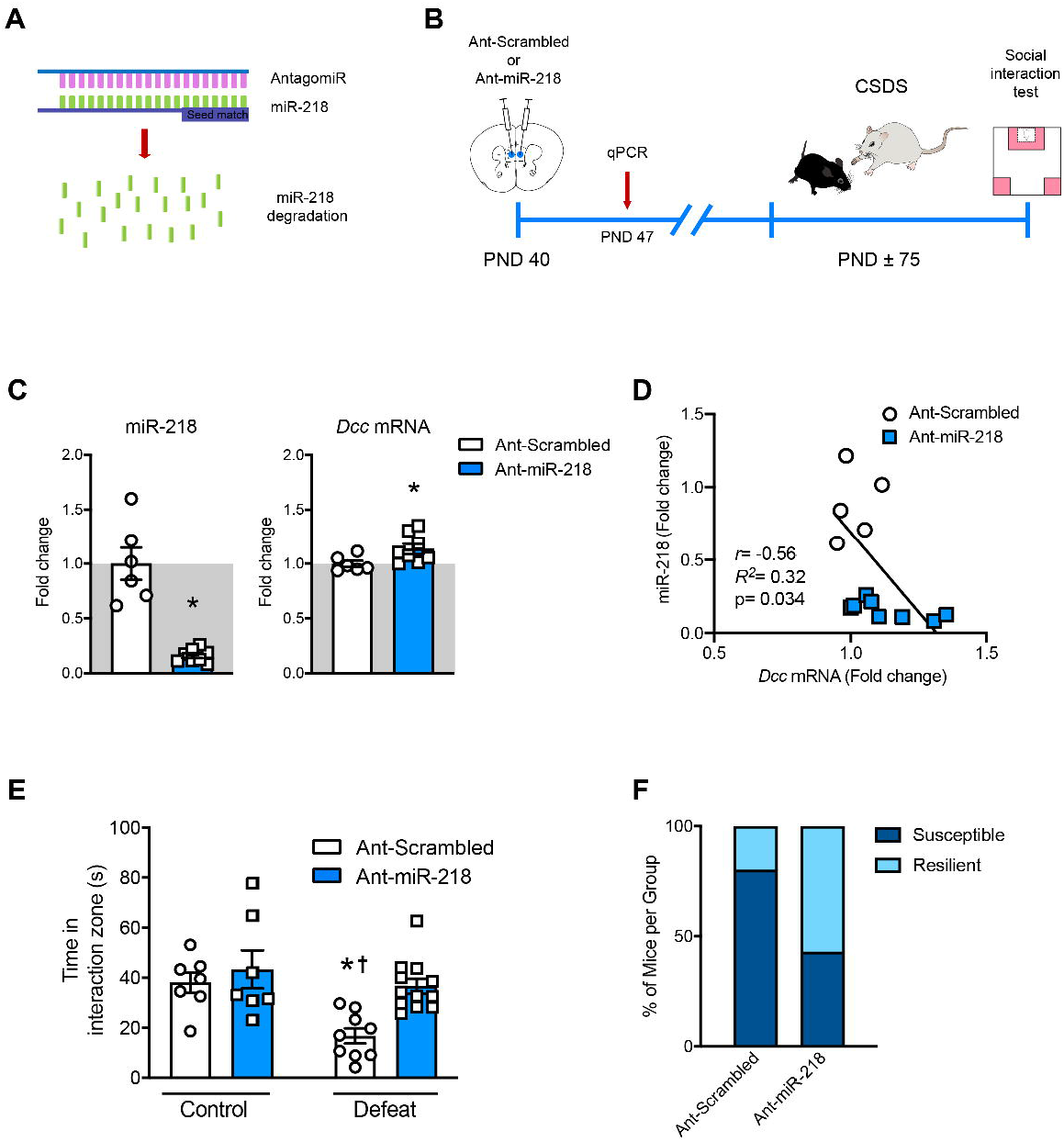
Downregulation of miR-218 in adolescence potentially promotes resilience to stress in adulthood. **(a)** Schematic illustration of the miR-218 antagomir (Ant-miR-218). **(b)** Timeline of experiment. **(c)** Significant downregulation of miR-218 and upregulation of *Dcc* mRNA in the mPFC following Ant-miR-218 infusion. miR-218: t_(12)_=6.49; *p<0.0001, different from Ant-scrambled; *Dcc* mRNA: t_(12)_=2.28; *p=0.041, different from Ant-scrambled. **(d)** Negative correlation of miR-218 and *Dcc* mRNA. **(e)** Time in the social interaction zone with target present: Two-way ANOVA: main effect of stress: F_(1,31)_= 10.10; p=0.0033; main effect of infusion: F_(1,31)_= 8.26; p= 0.0077, and stress by infusion interaction: F_(1,31)_= 2.79; p=0.1). Post hoc Tukey test shows reduced time in the social interaction zone in Ant-scrambled-Defeat in comparison to Ant-scrambled-Control, *p<0.05; Ant-miR-218-Control, †p<0.01 and Ant-miR-218-Defeat, †p<0.01.

Tissue from a small subset of mice was collected one week after surgery for antagomir validation (**FIGURE 3C,D)**. As expected, adolescent mice that received microinfusions of Ant-miR-218 exhibit decreased miR-218 expression (~70%) in the mPFC in comparison to their Ant-Scrambled counterpart. Furthermore, there is a slight but significant increase in *Dcc* mRNA expression (~20%) in mice infused with Ant-miR-218 (**FIGURE 3C**). Interestingly, the levels of miR-218 and *Dcc* mRNA in adolescence also correlate in a negative manner, consistent with our previous reports(30, 46).

Mice infused with Ant-scrambled during adolescence and exposed to CSDS in adulthood, exhibit a significant reduction in the time spent in the interaction zone when the CD-1 target is present, in comparison to non-stressed control groups. Notably, this effect is prevented in adult mice that received Ant-miR-218 infusions in the mPFC in adolescence (**FIGURE 3E**). Indeed, infusion of Ant-miR-218 in adolescence increases the proportion of “resilient” mice in comparison to Ant-scrambled infusion (**FIGURE 3F**). This suggests that reducing miR-218 expression in the mPFC specifically during adolescence can protect against vulnerability to adult exposure to stress.

## DISCUSSION

Discovering biomarkers during adolescence, a very sensitive time window for the onset of MDD, is urgently needed. It is also a critical step required for the implementation and/or modification of existing preventive and intervention strategies directed at those young individuals who are at higher risk of developing depression. Previous human research measured the unbiased transcriptional profile in the blood or hormone panels of children and adolescents with MDD with or without experiences of negative environmental factors, such as early life trauma(26, 52, 53). Genes related to the stress response or inflammation, including *NR3C1, TNF, TNFR1* and *IL1B*, have been significantly linked to symptoms of MDD(26, 52, 53), whereas DNA methylation of genes associated with neurodevelopment have been related to alterations in brain connectivity and early depressive symptoms(54, 55). In adult human studies, profiling of miRNAs using microarrays or small RNA sequencing have identified a few miRNAs as potential biomarkers of MDD, including miR-1202, miR-135, miR-146a/b-5, miR-425-3p, miR-24-3p, miR-941, and miR-589, which have also been found to be altered in postmortem brain tissue and plasma or blood-derived cells of MDD subjects (for reviews see(23, 24, 28, 29, 56–58)). These correlational studies open the possibility to conduct a systematic evaluation of these potential biomarkers across the lifespan to understand their functional implication in MDD vulnerability and how their alterations in the brain can be detected in blood using preclinical research strategies.

In this study we used rodent models to show that the levels of miR-218 in the mPFC and in blood change in parallel across development with a significant increase from early adolescence to adulthood. This finding is consistent with our previous observation showing that bidirectional manipulation of miR-218 in the adult mPFC associates with corresponding changes of circulating miR-218(31), and supports our idea that stress-induced alterations of miR-218 the mPFC can be observed in blood. The fact that high levels of circulating miR-218 of *adolescent stress-naïve mice* are associated with susceptibility to CSDS in adulthood suggests that an accelerated developmental expression of miR-218 could render this population more vulnerable to future stress. Indeed, analysis of blood expression of miR-218 after adult CSDS revealed that only control and resilient mice exhibit the expected developmental increase of circulating miR-218 levels; while susceptible mice show a reduction in miR-218 levels as we had observed before(31). These findings could indicate that mice that show susceptibility to adult CSDS already display “adult-like levels” of miR-218 in blood during adolescence, failing to maintain optimal levels after stress exposure. To our knowledge this is the first study reporting circulating expression of a miRNA as a potential developmental biomarker to stress-related disorders, including depression. Our results also show that the regulation of miR-218 by environmental factors is highly dependent on the specific critical period of development.

A question that remains to be answered relates to the different roles of miR-218 across development in the mPFC and how its manipulation leads to opposite effects in adolescent versus adulthood. Experimental downregulation of miR-218 during *adolescence* prevents susceptibility to adult exposure to CSDS (**FIGURE 3E,F**), whereas reducing the levels of miR-218 in the *adult* mPFC induces stress vulnerability(31). Interestingly, overexpression of miR-218 protects against CSDS-induced morphological alterations by increasing the density of thin spines. This suggests that miR-218 in the adult mPFC facilitates the formation of thin dendritic spines on pyramidal neurons under stressful conditions, and that these changes in spine plasticity might be protective for the mature brain. Evidence shows that miRNAs can regulate gene expression with high temporal and spatial specificity to mediate processes such as neuronal cell differentiation and maturation, neurogenesis, synapse formation and synaptic plasticity(59–61) and their function is tailored to selective developmental stages(62). Because miR-218 appears to maintain optimal levels of *Dcc* mRNA in the mPFC throughout life (**Fig. 1cd**), increased levels of miR-218 during adolescence, a period where its levels are transitioning from low-to-high, are likely to impair DCC’s normative developmental expression. This is likely to result in altered mPFC synapse formation, priming the brain to be overreactive to adverse experiences(63, 64). Future studies are needed to understand the changes in mPFC circuitry as the result of manipulating miR-218 in adolescence.

Beyond their potential role as biomarkers, miRNAs can be functionally involved in the etiology of a specific disease and, consequently, serve as promising targets for miRNA-based drug development(65, 66). Indeed, different strategies have been developed to manipulate, locally or peripherally, the levels of specific miRNAs by the use of mimics or antagomirs(66). Recent evidence demonstrates that intranasal delivery is a noninvasive method that allows some drugs to bypass the blood–brain barrier and rapidly reach the CNS(67, 68). This method has been successfully used in humans to treat psychiatric disorders, including schizophrenia(69) or depression(70). For example, intranasal administration of ketamine exerts rapid antidepressant effects in individuals suffering from treatment resistant depression(70–72). Furthermore, intranasal delivery of antagomirs has been proven to influence depression-like behaviors in adult mice exposed to CSDS(31, 73). Our previous finding demonstrating that intranasal administration of Ant-miR-218 turns adult resilient mice into susceptible, when evaluated in a second social interaction test(31), opens the possibility of manipulating miR-218 non-invasively and with precise temporal resolution. Delivering Ant-miR-218 intranasally during adolescence may have an enduring prophylactic antidepressant effect.

## ACKNOWLEDGMENTS AND DISCLOSURES

This work was funded by Canadian Institute for Health Research (C.F. MOP-74709), the National Institute on Drug Abuse (C.F. R01DA037911), the Natural Science and Engineering Research Council of Canada (C.F. 2982226) and the National Institute of Mental Health (E.J.N. P50MH096890 and R01MH051399). C.F. is a research scholar of the Fonds de Recherche du Québec - Santé. A.T.B. received the Integrated Program in Neuroscience fellowship. We thank Carlos Torres-Berrío for help in the graphic design.

The authors declare no potential conflict of interest.

## CONTRIBUTIONS

A.T.B. and C.F. conceived and designed the project. A.T.B., A.M., M.G., and S.C., performed gene expression experiments with mouse brain tissue, stereotaxic surgeries, antagomiR infusions, and behavioral experiments. E.J.N. provided reagents and substantial contribution to the interpretation of data. A.T.B and C.F. wrote the manuscript. A.T.B., C.F., A.M., and E.J.N. discussed the results, and reviewed and edited the final manuscript.

## Notes

### Competing Interest Statement

The authors have declared no competing interest.

## REFERENCES

1. Birmaher B, Brent D (2007): Practice Parameter for the Assessment and Treatment of Children and Adolescents With Depressive Disorders. Journal of the American Academy of Child & Adolescent Psychiatry. 46:1503–1526.

2. Davey CG, Yücel M, Allen NB (2008): The emergence of depression in adolescence: Development of the prefrontal cortex and the representation of reward. Neuroscience & Biobehavioral Reviews. 32:1–19.

3. Hammen C (2009): Adolescent Depression: Stressful Interpersonal Contexts and Risk for Recurrence. Current directions in psychological science. 18:200–204.

4. Brenhouse HC, Andersen SL (2011): Developmental trajectories during adolescence in males and females: a cross-species understanding of underlying brain changes. Neuroscience and biobehavioral reviews. 35:1687–1703.

5. Auerbach RP, Admon R, Pizzagalli DA (2014): Adolescent Depression: Stress and Reward Dysfunction. Harvard review of psychiatry. 22:139–148.

6. Gobinath AR, Mahmoud R, Galea LAM (2015): Influence of sex and stress exposure across the lifespan on endophenotypes of depression: focus on behavior, glucocorticoids, and hippocampus. Frontiers in Neuroscience. 8.

7. Kessler RC, Amminger GP, Aguilar-Gaxiola S, Alonso J, Lee S, Ustun TB (2007): Age of onset of mental disorders: A review of recent literature. Current opinion in psychiatry. 20:359–364.

8. LeMoult J, Humphreys KL, Tracy A, Hoffmeister J-A, Ip E, Gotlib IH (2019): Meta-analysis: Exposure to Early Life Stress and Risk for Depression in Childhood and Adolescence. Journal of the American Academy of Child & Adolescent Psychiatry.

9. Association AP (2013): Diagnostic and Statistical Manual of Mental Disorders - DSM–5. 5th ed. ed.: American Psychiatric Association, Washington, DC.

10. Shore L, Toumbourou JW, Lewis AJ, Kremer P (2018): Review: Longitudinal trajectories of child and adolescent depressive symptoms and their predictors – a systematic review and meta-analysis. Child and Adolescent Mental Health. 23:107–120.

11. Torres-Berrío A, Issler O, Parise EM, Nestler EJ (2019): Unraveling the epigenetic landscape of depression: focus on early life stress. Dialogues Clin Neurosci. 21:341–357.

12. Akil H, Gordon J, Hen R, Javitch J, Mayberg H, McEwen B, et al. (2018): Treatment resistant depression: A multi-scale, systems biology approach. Neuroscience & Biobehavioral Reviews. 84:272–288.

13. Shirtcliff EA, Allison AL, Armstrong JM, Slattery MJ, Kalin NH, Essex MJ (2012): Longitudinal stability and developmental properties of salivary cortisol levels and circadian rhythms from childhood to adolescence. Developmental Psychobiology. 54:493–502.

14. Owens M, Herbert J, Jones PB, Sahakian BJ, Wilkinson PO, Dunn VJ, et al. (2014): Elevated morning cortisol is a stratified population-level biomarker for major depression in boys only with high depressive symptoms. Proceedings of the National Academy of Sciences. 111:3638–3643.

15. Khandaker GM, Stochl J, Zammit S, Goodyer I, Lewis G, Jones PB (2018): Childhood inflammatory markers and intelligence as predictors of subsequent persistent depressive symptoms: a longitudinal cohort study. Psychological Medicine. 48:1514–1522.

16. Chu AL, Stochl J, Lewis G, Zammit S, Jones PB, Khandaker GM (2019): Longitudinal association between inflammatory markers and specific symptoms of depression in a prospective birth cohort. Brain, Behavior, and Immunity. 76:74–81.

17. Iob E, Kirschbaum C, Steptoe A (2019): Persistent depressive symptoms, HPA-axis hyperactivity, and inflammation: the role of cognitive-affective and somatic symptoms. Mol Psychiatry.

18. Colich NL, Kircanski K, Foland-Ross LC, Gotlib IH (2015): HPA-axis reactivity interacts with stage of pubertal development to predict the onset of depression. Psychoneuroendocrinology. 55:94–101.

19. Humphreys KL, Moore SR, Davis EG, MacIsaac JL, Lin DTS, Kobor MS, et al. (2019): DNA methylation of HPA-axis genes and the onset of major depressive disorder in adolescent girls: a prospective analysis. Transl Psychiatry. 9:245.

20. King LS, Colich NL, LeMoult J, Humphreys KL, Ordaz SJ, Price AN, et al. (2017): The impact of the severity of early life stress on diurnal cortisol: The role of puberty. Psychoneuroendocrinology. 77:68–74.

21. LeMoult J, Ordaz SJ, Kircanski K, Singh MK, Gotlib IH (2015): Predicting first onset of depression in young girls: Interaction of diurnal cortisol and negative life events. Journal of Abnormal Psychology. 124:850–859.

22. Kentner AC, Cryan JF, Brummelte S (2019): Resilience priming: Translational models for understanding resiliency and adaptation to early life adversity. Developmental Psychobiology. 61:350–375.

23. Issler O, Haramati S, Paul Evan D, Maeno H, Navon I, Zwang R, et al. (2014): MicroRNA 135 Is Essential for Chronic Stress Resiliency, Antidepressant Efficacy, and Intact Serotonergic Activity. Neuron. 83:344–360.

24. Belzeaux R, Lin R, Turecki G (2017): Potential Use of MicroRNA for Monitoring Therapeutic Response to Antidepressants. CNS Drugs. 31:253–262.

25. Allen L, Dwivedi Y (2020): MicroRNA mediators of early life stress vulnerability to depression and suicidal behavior. Mol Psychiatry. 25:308–320.

26. Pajer K, Andrus BM, Gardner W, Lourie A, Strange B, Campo J, et al. (2012): Discovery of blood transcriptomic markers for depression in animal models and pilot validation in subjects with early-onset major depression. Transl Psychiatry. 2:e101–e101.

27. Chiang JJ, Cole SW, Bower JE, Irwin MR, Taylor SE, Arevalo J, et al. (2019): Depressive symptoms and immune transcriptional profiles in late adolescents. Brain, Behavior, and Immunity. 80:163–169.

28. Lopez JP, Lim R, Cruceanu C, Crapper L, Fasano C, Labonte B, et al. (2014): miR-1202 is a primate-specific and brain-enriched microRNA involved in major depression and antidepressant treatment. Nature Medicine. 20:764–768.

29. Lopez JP, Fiori LM, Cruceanu C, Lin R, Labonte B, Cates HM, et al. (2017): MicroRNAs 146a/b-5 and 425-3p and 24-3p are markers of antidepressant response and regulate MAPK/Wnt-system genes. 8:15497.

30. Torres-Berrío A, Lopez JP, Bagot RC, Nouel D, Dal Bo G, Cuesta S, et al. (2017): DCC Confers Susceptibility to Depression-like Behaviors in Humans and Mice and Is Regulated by miR-218. Biological Psychiatry. 81:306–315.

31. Torres-Berrío A, Nouel D, Cuesta S, Parise EM, Restrepo-Lozano JM, Larochelle P, et al. (2019): MiR-218: a molecular switch and potential biomarker of susceptibility to stress. Mol Psychiatry. 25:951–964.

32. Manitt C, Eng C, Pokinko M, Ryan RT, Torres-Berrio A, Lopez JP, et al. (2013): dcc orchestrates the development of the prefrontal cortex during adolescence and is altered in psychiatric patients. Transl Psychiatry. 3:e338.

33. Reynolds LM, Pokinko M, Torres Berrío A, Cuesta S, Lambert LC, Del Cid Pellitero E, et al. (2017): DCC receptors drive prefrontal cortex maturation by determining dopamine axon targeting in adolescence. Biological Psychiatry. 83:181–192.

34. Hoops D, Flores C (2017): Making Dopamine Connections in Adolescence. Trends in Neurosciences. 40:709–719.

35. Dunn EC, Wiste A, Radmanesh F, Almli LM, Gogarten SM, Sofer T, et al. (2016): Genome-Wide Association Study (GWAS) and Genome-Wide by Environment Interaction Study (GWEIS) of Depressive Symptoms in African American and Hispanic/Latina Women. Depression and Anxiety. 33:265–280.

36. Ward J, Strawbridge RJ, Bailey MES, Graham N, Ferguson A, Lyall DM, et al. (2017): Genome-wide analysis in UK Biobank identifies four loci associated with mood instability and genetic correlation with major depressive disorder, anxiety disorder and schizophrenia. Transl Psychiatry. 7:1264.

37. Zeng Y, Navarro P, Fernandez-Pujals AM, Hall LS, Clarke T-K, Thomson PA, et al. (2017): A Combined Pathway and Regional Heritability Analysis Indicates NETRIN1 Pathway Is Associated With Major Depressive Disorder. Biological Psychiatry. 81:336–346.

38. Aberg KA, Shabalin AA, Chan RF, Zhao M, Kumar G, van Grootheest G, et al. (2018): Convergence of evidence from a methylome-wide CpG-SNP association study and GWAS of major depressive disorder. Transl Psychiatry. 8:162.

39. Elliott LT, Sharp K, Alfaro-Almagro F, Shi S, Miller KL, Douaud G, et al. (2018): Genome-wide association studies of brain imaging phenotypes in UK Biobank. Nature. 562:210–216.

40. Arnau-Soler A, Macdonald-Dunlop E, Adams MJ, Clarke T-K, MacIntyre DJ, Milburn K, et al. (2019): Genome-wide by environment interaction studies of depressive symptoms and psychosocial stress in UK Biobank and Generation Scotland. Transl Psychiatry. 9:14.

41. Kichaev G, Bhatia G, Loh P-R, Gazal S, Burch K, Freund MK, et al. (2019): Leveraging Polygenic Functional Enrichment to Improve GWAS Power. The American Journal of Human Genetics. 104:65–75.

42. Barbu MC, Zeng Y, Shen X, Cox SR, Clarke T-K, Gibson J, et al. (2019): Association of Whole-Genome and NETRIN1 Signaling Pathway–Derived Polygenic Risk Scores for Major Depressive Disorder and White Matter Microstructure in the UK Biobank. Biological Psychiatry: Cognitive Neuroscience and Neuroimaging. 4:91–100.

43. Lee PH, Anttila V, Won H, Feng Y-CA, Rosenthal J, Zhu Z, et al. (2019): Genomic Relationships, Novel Loci, and Pleiotropic Mechanisms across Eight Psychiatric Disorders. Cell. 179:1469–1482.e1411.

44. Vosberg DE, Leyton M, Flores C (2020): The Netrin-1/DCC guidance system: dopamine pathway maturation and psychiatric disorders emerging in adolescence. Mol Psychiatry. 25:297–307.

45. Torres-Berrío A, Hernandez G, Nestler EJ, Flores C (2020): The Netrin-1/DCC guidance cue pathway as a molecular target in depression: Translational evidence. Biological Psychiatry.

46. Cuesta S, Restrepo-Lozano JM, Silvestrin S, Nouel D, Torres-Berrío A, Reynolds LM, et al. (2017): Non-Contingent Exposure to Amphetamine in Adolescence Recruits miR-218 to Regulate Dcc Expression in the VTA. Neuropsychopharmacology. 43:900–911.

47. Paxinos G, Franklin K (2013): Paxinos and Franklin’s the Mouse Brain in Stereotaxic Coordinates. Boston, Amsterdam: Elsevier/Academic Press.

48. Goldman JS, Ashour MA, Magdesian MH, Tritsch NX, Harris SN, Christofi N (2013): Netrin-1 promotes excitatory synaptogenesis between cortical neurons by initiating synapse assembly. J Neurosci. 33.

49. Reynolds LM, Yetnikoff L, Pokinko M, Wodzinski M, Epelbaum JG, Lambert LC, et al. (2018): Early Adolescence is a Critical Period for the Maturation of Inhibitory Behavior. Cerebral Cortex. 29:3676–3686.

50. Small EM, Sutherland LB, Rajagopalan KN, Wang S, Olson EN (2010): MicroRNA-218 Regulates Vascular Patterning by Modulation of Slit-Robo Signaling. Circulation Research. 107:1336–1344.

51. Stein E, Tessier-Lavigne M (2001): Hierarchical Organization of Guidance Receptors: Silencing of Netrin Attraction by Slit Through a Robo/DCC Receptor Complex. Science. 291:1928–1938.

52. Minelli A, Magri C, Giacopuzzi E, Gennarelli M (2018): The effect of childhood trauma on blood transcriptome expression in major depressive disorder. Journal of Psychiatric Research. 104:50–54.

53. Spindola LM, Pan PM, Moretti PN, Ota VK, Santoro ML, Cogo-Moreira H, et al. (2017): Gene expression in blood of children and adolescents: Mediation between childhood maltreatment and major depressive disorder. Journal of Psychiatric Research. 92:24–30.

54. Kaufman J, Wymbs NF, Montalvo-Ortiz JL, Orr C, Albaugh MD, Althoff R, et al. (2018): Methylation in OTX2 and related genes, maltreatment, and depression in children. Neuropsychopharmacology. 43:2204–2211.

55. de Araújo CM, Hudziak J, Crocetti D, Wymbs NF, Montalvo-Ortiz JL, Orr C, et al. (2020): Tubulin Polymerization Promoting Protein (TPPP) gene methylation and corpus callosum measures in maltreated children. Psychiatry Research: Neuroimaging. 298:111058.

56. Chen RJ, Kelly G, Sengupta A, Heydendael W, Nicholas B, Beltrami S, et al. (2015): MicroRNAs as biomarkers of resilience or vulnerability to stress. Neuroscience. 305:36–48.

57. Fries GR, Zhang W, Benevenuto D, Quevedo J (2019): MicroRNAs in Major Depressive Disorder. In: Guest PC, editor. Reviews on Biomarker Studies in Psychiatric and Neurodegenerative Disorders. Cham: Springer International Publishing, pp 175–190.

58. Tavakolizadeh J, Roshanaei K, Salmaninejad A, Yari R, Nahand JS, Sarkarizi HK, et al. (2018): MicroRNAs and exosomes in depression: Potential diagnostic biomarkers. Journal of Cellular Biochemistry, n/a-n/a.

59. Schratt GM, Tuebing F, Nigh EA, Kane CG, Sabatini ME, Kiebler M, et al. (2006): A brain-specific microRNA regulates dendritic spine development. Nature. 439:283–289.

60. Aksoy-Aksel A, Zampa F, Schratt G (2014): MicroRNAs and synaptic plasticity—a mutual relationship. Philosophical Transactions of the Royal Society B: Biological Sciences. 369.

61. Sambandan S, Akbalik G, Kochen L, Rinne J, Kahlstatt J, Glock C, et al. (2017): Activity-dependent spatially localized miRNA maturation in neuronal dendrites. Science. 355:634–637.

62. Alvarez-Garcia I, Miska EA (2005): MicroRNA functions in animal development and human disease. Development. 132:4653.

63. Siegel G, Obernosterer G, Fiore R, Oehmen M, Bicker S, Christensen M, et al. (2009): A functional screen implicates microRNA-138-dependent regulation of the depalmitoylation enzyme APT1 in dendritic spine morphogenesis. Nat Cell Biol. 11:705–716.

64. Rocchi A, Moretti D, Lignani G, Colombo E, Scholz-Starke J, Baldelli P, et al. (2019): Neurite-Enriched MicroRNA-218 Stimulates Translation of the GluA2 Subunit and Increases Excitatory Synaptic Strength. Molecular Neurobiology.

65. Sun P, Liu DZ, Jickling GC, Sharp FR, Yin K-J (2018): MicroRNA-based therapeutics in central nervous system injuries. Journal of Cerebral Blood Flow & Metabolism. 38:1125–1148.

66. Schmidt MF (2014): Drug target miRNAs: chances and challenges. Trends in Biotechnology. 32:578–585.

67. Dhuria SV, Hanson LR, Frey WH (2010): Intranasal delivery to the central nervous system: Mechanisms and experimental considerations. Journal of Pharmaceutical Sciences. 99:1654–1673.

68. Hanson LR, Fine JM, Svitak AL, Faltesek KA (2013): Intranasal Administration of CNS Therapeutics to Awake Mice. JoVE.e4440.

69. Katare YK, Piazza JE, Bhandari J, Daya RP, Akilan K, Simpson MJ, et al. (2017): Intranasal delivery of antipsychotic drugs. Schizophrenia Research. 184:2–13.

70. Quintana DS, Steen NE, Andreassen OA (2018): The Promise of Intranasal Esketamine as a Novel and Effective Antidepressant. JAMA Psychiatry. 75:123–124.

71. Krystal JH, Charney DS, Duman RS (2020): A New Rapid-Acting Antidepressant. Cell. 181:7.

72. Lapidus KAB, Levitch CF, Perez AM, Brallier JW, Parides MK, Soleimani L, et al. (2014): A Randomized Controlled Trial of Intranasal Ketamine in Major Depressive Disorder. Biological Psychiatry. 76:970–976.

73. Deng Z-F, Zheng H-L, Chen J-G, Luo Y, Xu J-F, Zhao G, et al. (2018): miR-214-3p Targets β-Catenin to Regulate Depressive-like Behaviors Induced by Chronic Social Defeat Stress in Mice. Cerebral Cortex.

